# The invasive land flatworm *Obama nungara* in La Réunion, a French island in the Indian Ocean, the first report of the species for Africa

**DOI:** 10.1101/2022.02.14.480416

**Authors:** Jean-Lou Justine, Amandine Delphine Marie, Romain Gastineau, Yoan Fourcade, Leigh Winsor

## Abstract

The land flatworm *Obama nungara*, a species originating from South America and already invasive in many European countries, is recorded from La Réunion, a French island in the Indian Ocean. This is the first record of *O. nungara* from this locality and also the first record of the species for Africa. Two specimens were collected, one from Petite France (commune of Saint Paul) and one from La Plaine des Grègues (commune of Saint Joseph); the two localities are widely separated, one in the Western part and one in the South-eastern part of the island. This suggests that the species is already present in several locations in La Réunion. The sightings were communicated to us in 2021, but it is likely that the species is already present since 2020. A molecular analysis of the specimen from Petite France showed that it had the same *cox1* haplotype as specimens previously recorded from several countries of Europe; it is hypothesized that the species was imported from Europe, probably from France. We mapped climatic suitability of the species in La Réunion and found that *O. nungara* could potentially invade a large part of the island. One record was apparently associated with the transport of plates of travertine, a construction material which has numerous cavities, suitable for the transport and survival of adult or cocoons of land flatworms.

## Introduction

Invasive species are permeating every part of the planet, even the most remote sites. Land flatworms are representative of this trend (Sluys, 2016). The New Guinea flatworm, *Platydemus manokwari* de Beauchamp, 1963, has been included in the list of the “100 world’s worst invasive alien species” (Lowe et al., 2000); it is present in many territories in the Pacific (Justine et al., 2014), has recently invaded continental USA (Justine et al., 2015), and is associated with the decline of native snails (Gerlach et al., 2020). Other examples include the hammerhead flatworms, especially *Bipalium kewense* Moseley, 1878, now in dozens of countries, or *Bipalium vagum* Jones & Sterrer, 2005, now widespread in many tropical countries (Justine et al., 2018; Winsor, 1983).

*Obama nungara* Carbayo et al., 2016 was recently described from a series of specimens, both from their land of origin, South America, and from Europe (Carbayo et al., 2016). It has now invaded several countries in Europe (Justine et al., 2020b). Molecular studies have shown that the species has three *cox1* haplotypes, designated as “Argentina 1”, “Argentina 2” and “Brazil” according to their region of origin. The Brazil haplotype has never been found outside of Brazil; the Argentina 2 haplotype has been recorded only in Argentina and a few localities in Spain; in contrast, the Argentina 1 haplotype has been found in several countries in Europe, including France, Spain, Portugal, Italy, Switzerland and UK (Justine et al., 2020b).

*Obama nungara* is a predator of earthworms and molluscs and thus is a potential threat for the biodiversity of soils. It is able to proliferate at high rates in gardens in temperate climates, with thousands of specimens in a single garden (Justine et al., 2020b).

In this paper, we report the first finding of *O. nungara* in La Réunion, a French island in the Indian Ocean. We show that the *cox1* haplotype of this specimen is compatible with an importation from Europe, probably metropolitan France. In addition, we mapped climatic suitability of the species in La Réunion, projected from a previously published species distribution model (Fourcade, 2021), showing that this species could invade a large part of the island.

## Material and Methods

### Collection of specimens

Citizen science is highly effective for collecting information of land flatworms (Justine et al., 2022; Justine et al., 2020a; Justine et al., 2015; Justine et al., 2014, 2018, 2020b; Mori et al., 2022). One of us (ADM) initiated a citizen science project in La Réunion about land flatworms, supported by the local association ARBRE (Agence de Recherche pour la Biodiversité à La Réunion *https://arb-reunion.fr/*). Among the many records obtained from participating individuals, a few flatworms were morphologically similar to *Obama nungara*. Photographs of the living flatworms were taken, and a few specimens of flatworms were collected. The specimen from Petite France was registered in the collections of the Muséum national d’Histoire naturelle under number MNHN JL449, and then a part of the body was taken for molecular study and sent to Poland (RG). The specimen from la Plaine des Grègues was registered as MNHN JL452 and no molecular study was undertaken.

### Molecular methods: sequencing, alien DNA, and haplotype network

The complete mitogenome of specimen JL449 was sequenced in the context of a project in progress. Sequencing was performed following the exact same protocol as described in detail in a recent paper (Justine et al., 2022), with a k-mer of 125. The sequence corresponding to the *cox1* gene was identified using ORF finder on the NCBI portal (https://www.ncbi.nlm.nih.gov/orffinder) and by comparison with a published complete mitogenome (Solà et al., 2015). The *cox1* sequence was deposited in GenBank as OM451228.

The pool of contigs obtained after assembly was also datamined to look for the presence of alien DNA following Justine et al. (2022). The phylogenetic tree (not shown) obtained on this contaminant DNA was based on the partial 5.8S and ITS1, which were aligned with corresponding sequences of slugs (Quinteiro et al., 2005; Rowson et al., 2014), trimmed and computed for a Maximum Likelihood (ML) phylogeny with the General Time Reverse model and 100 bootstrap replicates, all steps performed by MEGAX (Kumar et al., 2018).

We constructed an haplotype network using a matrix made public (Justine et al., 2020b) and simply added one taxon (the *cox1* sequence of specimen JL449) and one trait label (“La Réunion”). In the nexus file, traits were the geographical origin of the species, with number of traits set to 10, and trait labels set as: Argentina, Brazil, UK (including Guernsey), Portugal, Spain, Italy, Switzerland, Belgium, Metropolitan France, and La Réunion. The matrix had 92 taxa and was 255 bases in length. We used PopART (Leigh & Bryant, 2015) to create the haplotype network; the method of Templeton, Crandall and Sing (“TCS”) was used to infer relationships among samples (Clement et al., 2002).

### Species distribution modelling

We produced a map of climatic suitability of O. nungara in La Réunion using an already published species distribution model (Fourcade, 2021). Briefly, the a MaxEnt model (Phillips et al., 2017) was fitted via the ENMeval R package (Kass et al., 2021) using georeferenced records of O. nungara both from the species’ native range and from Europe, as well as bioclimatic predictors at 2.5 arc-min resolution. Here, we obtained the same bioclimatic variables at a finer resolution (30 arc-sec) and used them to project model predictions in La Réunion. Full details of the modelling parameters are available in Fourcade et al. (2021), where the employed model is referred to as ‘fully tuned’.

## Results

As part of our citizen science initiative, we obtained two records of flatworms resembling *Obama nungara* from La Réunion.

- Régis Bretzner found several specimens of flatworms, in his house located in Petite France, a village belonging administratively to the commune of Saint Paul, in the western part of the island. The specimens, ranging from a few millimetres to 4 cm, were seen mainly in his bathroom. The first specimen was found on 10 June 2021 **(Figure 1)**, and the participant recalled seeing flatworms until the end of July, but not since (telephone interview in January 2022). The bathroom had been recently redone by with plates of travertine. The travertine had many holes and cavities; it was bought locally, its origin was unknown, but it is likely that the stones were imported from France.
- Renaud Hoarau collected and photographed a specimen on 22 April 2021 in the garden of his house in la Plaine des Grègues in the commune of Saint Joseph, in the eastern part of the island. The specimens were seen in his garden. He moved into this house in 2020 and remembers seeing flatworms since he first arrived, and continually, up to January 2022 when we interviewed him by telephone. He provided a photograph taken 22 April 2021 in his home **(Figure 2)** and a photograph taken 25 January 2022 close to his home (not shown).

**Figure 1.**
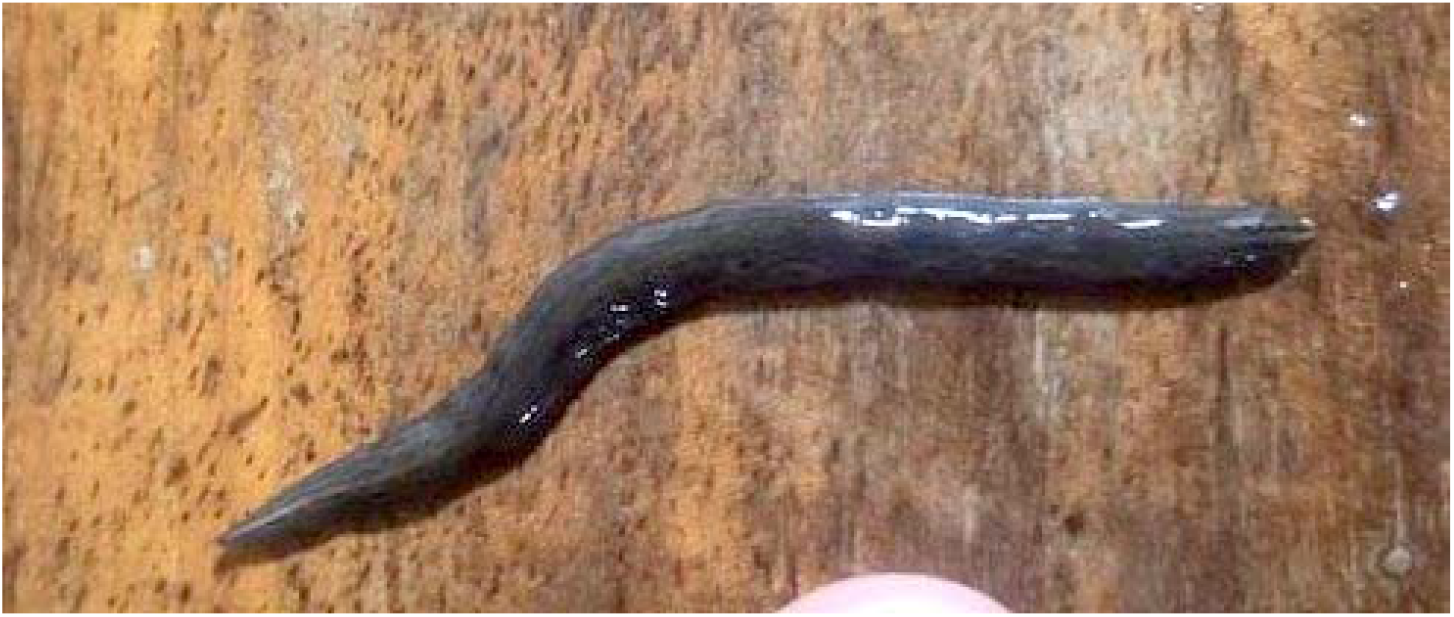
Photo of a live specimen of *Obama nungara*, taken in Petite France in the commune of Saint Paul, La Réunion. Photo by Régis Bretzner, taken 10 June 2021. The specimen is brown on the back, with darker brown striae, a reddish tip, and with a cream stripe running a short distance from the head towards the tail, but which soon disappears into the general colour of the back.

**Figure 2.**
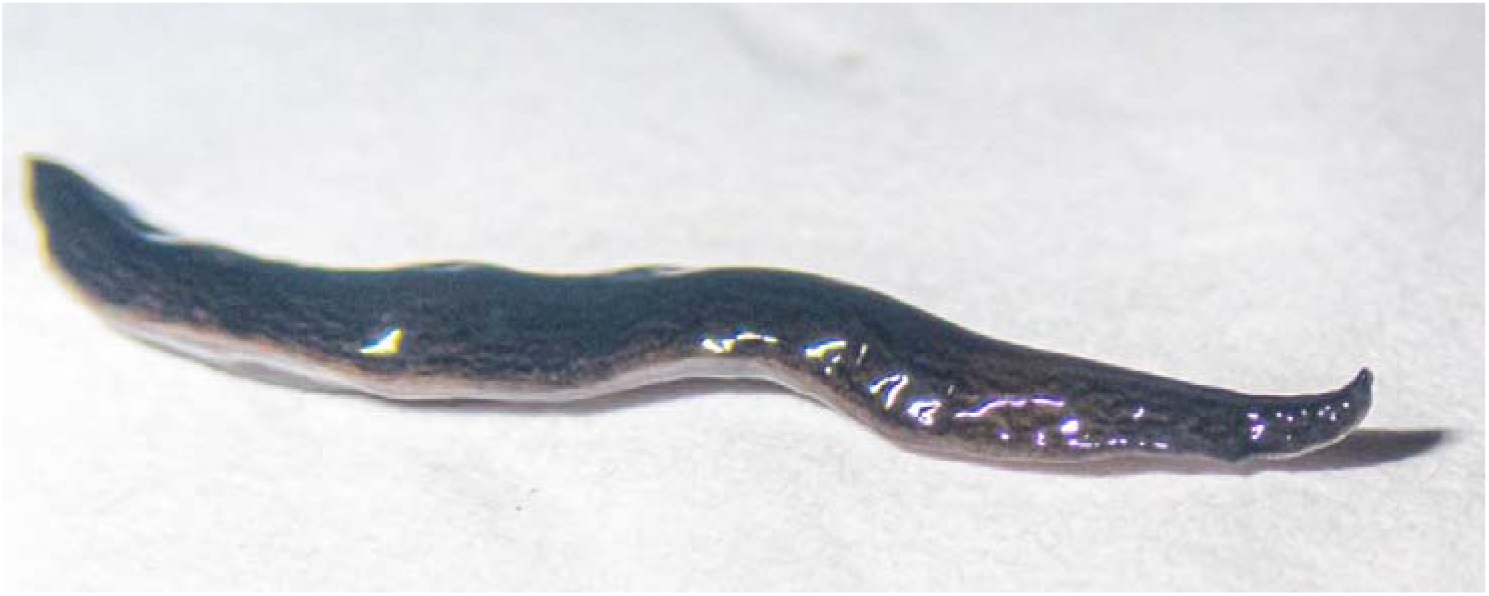
Photo of a live specimen of *Obama nungara*, taken in la Plaine des Grègues in the commune of Saint Joseph, La Réunion. Photo by Renaud Hoarau, taken 22 April 2021. The photo clearly shows the dark brown striae on the brownish coloured back, and the whitish under surface.

The living specimens on the photographs sent by both collectors showed the general characteristics of *Obama nungara*, with a brown dorsal colour with darker striae (**Figures 1-2**). The preserved specimens showed a green-brown dorsal colour and a beige ventral colour (not shown).

The complete *cox1* gene was 1737 bp long, which is identical to the reference from Solà et al. (2015) and has the same start and stop codons (GTG and TAA respectively). The haplotype network (**Figure 3**) showed that the specimen from La Réunion was identical to one of the haplotypes included within the Argentina 1 network. This haplotype includes samples from France, Spain, Argentina, Switzerland, and Belgium.

**Figure 3.**
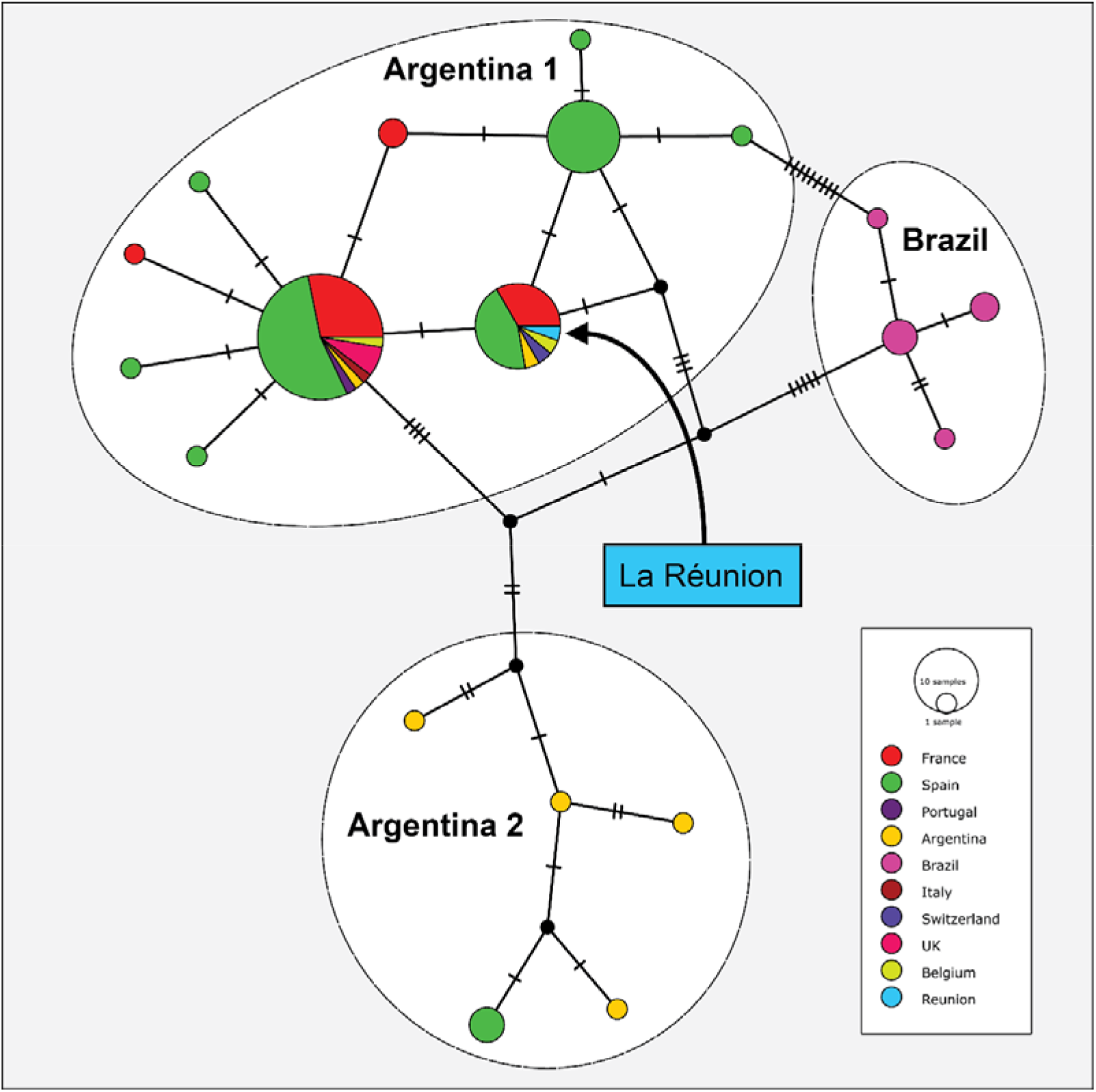
Haplotype network. The network was obtained by using a matrix made public (Justine et al., 2020b) and adding a single sequence, from the specimen MNHN JL449 from Petite France.

In our research of alien DNA, only few contigs with low coverage returned positive blastn results. Megablast queries showed that all these contigs belong to the cluster of nuclear ribosomal RNA genes of a slug of the genus *Arion*. The ML tree associates the alien DNA in a highly supported clade with *Arion intermedius* (Normand, 1852) isolate 56D (GenBank accession number AY316286.1), a specimen from Aran Valley, Lleida, Spain.

Modelling (**Figure 4**) showed that ca. 57% of the island of La Réunion was suitable for *O. nungara* (based on the threshold that maximises model sensitivity and specificity); in addition, about 38% of the island had a very high climatic suitability (> 0.8). Based on the analysis of the model’s response curves (Fourcade, 2021), it appears that the limiting factor was mainly the mean temperature of the warmest quarter (BIO 10 bioclimatic variable), coastal areas being too warm while high-altitude areas are too cold. It is noteworthy that both localities where *O. nungara* specimens were found are inside the zone with highest suitability (Plaine des Grègues: 0.92; Petite France: 0.89).

**Figure 4.**
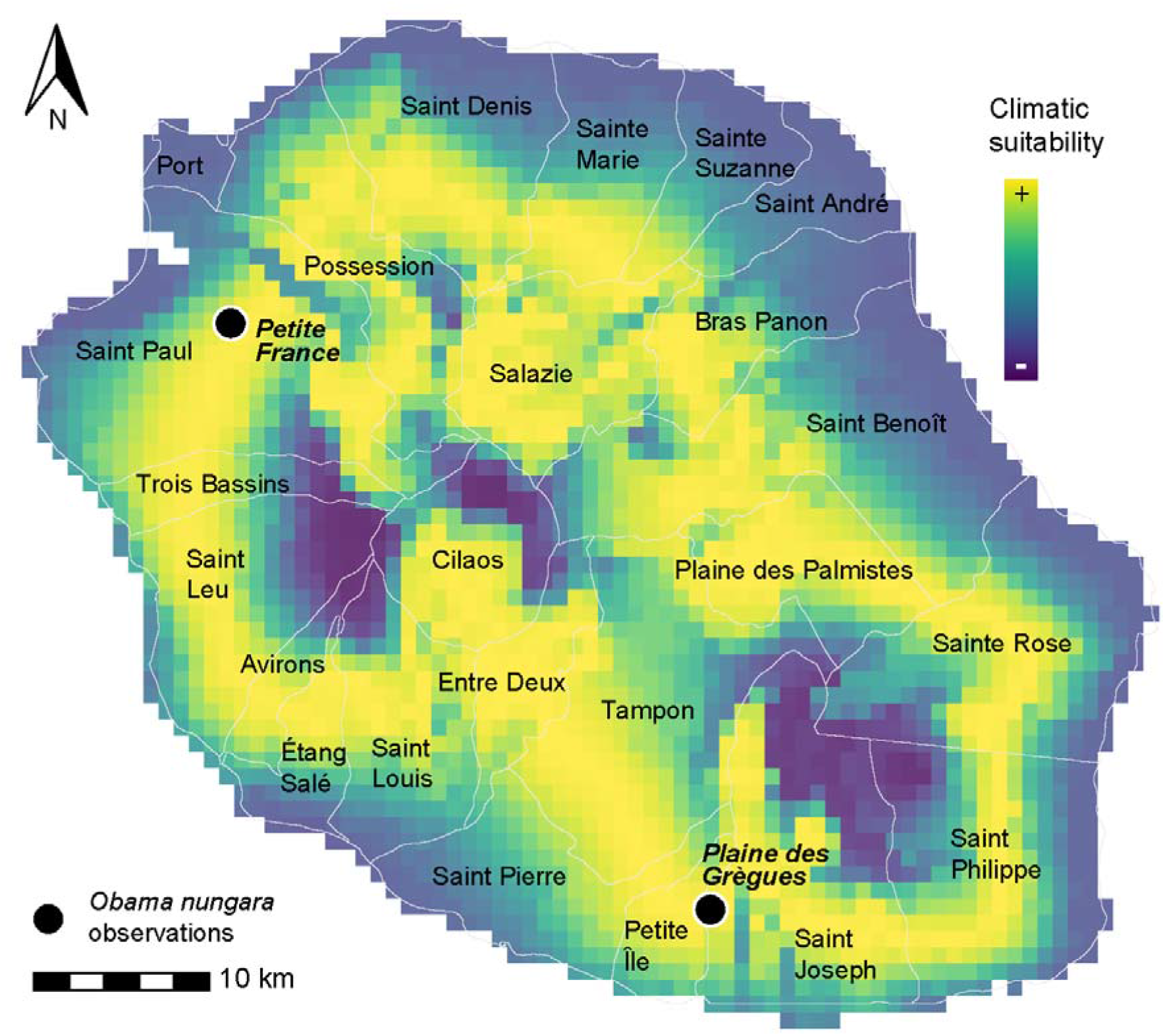
Climatic suitability of *Obama nungara* in La Réunion. Names of communes and their limits (white lines) are indicated. The two localities where the specimens were found (Petite France and Plaine des Grègues) are indicated.

## Discussion

*Obama nungara*, originating from South America (Brazil and Argentina) (Carbayo et al., 2016) has now invaded many countries in Europe. In 2020, a study found that Guernsey Island, UK, Spain, Portugal, France, Belgium, Italy, Ireland, Switzerland, and Madeira Island were already invaded (Justine et al., 2020b). More recently, the species was reported from Azores Islands (Lago-Barcia et al., 2020), The Netherlands (de Waart et al., 2021), and finally

Austria and Germany (Justine et al., unpublished) and Slovakia (Čapka & Čejka, 2021), therefore falsifying a previous statement (Justine et al., 2020b) that “no record was known from Germany or any country East of Germany”. There are also reports of specimens resembling *O. nungara* from USA, Mexico, Costa Rica, Bolivia, and Chile in iNaturalist (*www.iNaturalist.org*), but none of these citizen science records has been published nor confirmed by molecular methods.

The present record in the island of La Réunion, a French island located east of Madagascar in the Indian Ocean, is the first for this territory. As such, it is also the first report of *Obama nungara* for Africa. Another alien flatworm, *Bipalium vagum*, has been recorded in La Réunion (Justine et al., 2018). Our current citizen science initiative has shown that other flatworm species are present, and these records will be published in papers to come.

There are two studies on the potential distribution of *Obama nungara* worldwide. One used built-in parameters in the software (Negrete et al., 2020) and one used more elaborate methods (Fourcade, 2021). In this second study, it was shown that *O. nungara* had the potential to invade a large number of territories in the world. This includes a significant part of Africa, in particular south and east Africa, as well as Madagascar and its neighbouring islands. We were able, here, to specify at a finer scale the localities of La Réunion that appear climatically suitable for *O. nungara*. Model projections reveal that most of the areas of intermediate altitude have the potential to be invaded, because they harbour mild temperatures during the hot season that are needed for *O. nungara* survival. A more precise assessment of invasion risk would require a finer characterisation of *O. nungara*’s ecological niche, e.g. with regard to microclimate, soil properties, as well as biological interactions with its prey and predators.

The present report is based on only two records, but we have no doubt that these are only the visible part of the iceberg. In the two islands of Martinique and Guadeloupe (Antilles) a similar situation was found for the New Guinea flatworm, *Platydemus manokwari*. Only two years after the first report (Justine & Winsor, 2020), it was found that the flatworm had already invaded most of Guadeloupe (Justine et al., 2021). Similarly, the invasive flatworm *Amaga expatria* was found in many locations in the two islands (Justine et al., 2020a). This is again an illustration that islands are fragile ecosystems which are more sensitive to invasion by alien species. We fear that the same will happen in La Réunion with *O. nungara* (and possibly other flatworm species).

Since the feeding behaviour of *O. nungara* has not been yet directly observed in la Réunion, the results we obtained on the alien DNA associated to the specimen have to be taken with care, keeping in mind that they concern a single specimen. Detecting a European species, the Hedgehog Slug *Arion intermedius* as prey, was unexpected. However, this species is a known introduction in many countries and territories, including La Réunion (Auffenberg et al., 2021).

In the case of the invasion of La Réunion by *O. nungara*, it is noteworthy that the specimens most probably do not come from their land of origin (Brazil or Argentina) but from another continent (Europe) and, because of the high level of commerce between La Réunion and France, probably from the latter.

Land flatworms are usually considered to be spread internationally by the transport of plants (Murchie & Justine, 2021; Sluys, 2016). Indeed, adults flatworms can survive in the earth of potted plants, and cocoons can also go undetected in the same situation. Our current study suggests that the transport of travertine stone was implicated in the possible spread of *O. nungara*. We do not know whether the flatworms were directly imported from Europe to La Réunion in a stack of travertine stone, or if following delivery to La Réunion, the stack was colonized by a local proliferation of the flatworm in an area of the island where the stone was kept before being sold to the public; the finding of many specimens by the observer in Saint Joseph favours the second hypothesis. In both cases, this hypothesis of transport of the flatworms together with the stone deserves consideration. Indeed, decorative stone such as travertine is often transported as stacks with the plates separated from each other by very thin spaces. Considering that travertine offers many spaces and cavities, which can harbour either adults or cocoons, this stone offers ideal conditions for the transport of flatworms such as *Obama nungara*. Transport of inert materials such as flat slices or tiles of stone should be considered a possible way of transport for invasive flatworms, in addition to the transport of plants.

## Acknowledgements

The thank the participants in our citizen science project.

## References

Auffenberg, K., Bank, R., Bieler, R., Bouchet, P., Faber, M., Finn, J., et al. (2021) Arion intermedius Normand, 1852. In: Bánki (Ed), MolluscaBase ver. (10/2021).

Čapka, J. & Čejka, T. (2021) First record of Obama nungara in Slovakia (Platyhelminthes: Geoplanidae). Biodiversity & Environment, Vol. 13, No. 2, 13, 41-44.

Carbayo, F., Alvarez-Presas, M., Jones, H. D. & Riutort, M. (2016) The true identity of Obama (Platyhelminthes: Geoplanidae) flatworm spreading across Europe. Zoological Journal of the Linnean Society, 177, 5–28. http://doi.org/10.1111/zoj.12358

Clement, M., Snell, Q., Walke, P., Posada, D. & Crandall, K. (2002) TCS: estimating gene genealogies. In: Proceedings of the 16th International Parallel and Distributed Process Symposium 2:184., Fort Lauderdale, Florida.

de Waart, S., Thunnissen, N. & Sluys, R. (2021) Exotische landplatwormen in Nederland (Platyhelminthes: Tricladida). Nederlandse Faunistische Mededelingen, 47, 15–27.

Fourcade, Y. (2021) Fine-tuning niche models matters in invasion ecology. A lesson from the land planarian Obama nungara. Ecological Modelling, 457, 109686. http://doi.org/10.1016/j.ecolmodel.2021.109686

Gerlach, J., Barker, G. M., Bick, C. S., Bouchet, P., Brodie, G., Christensen, C. C., et al. (2020) Negative impacts of invasive predators used as biological control agents against the pest snail Lissachatina fulica: the snail Euglandina ‘rosea’and the flatworm Platydemus manokwari. Biological Invasions, https://doi.org/10.1007/s10530-10020-02436-w.

Justine, J.-L., Gastineau, R., Gros, P., Ruzzier, E., Charles, L. & Winsor, L. (2022) Hammerhead flatworms (Platyhelminthes, Geoplanidae, Bipaliinae): mitochondrial genomes and description of two new species from France, Italy, and Mayotte. PeerJ, 10, e12725 http://doi.org/10.7717/peerj.12725

Justine, J.-L., Gey, D., Thévenot, J., Gastineau, R. & Jones, H. D. (2020a) The land flatworm Amaga expatria (Geoplanidae) in Guadeloupe and Martinique: new reports and molecular characterization including complete mitogenome. PeerJ, 8, e10098. http://doi.org/10.7717/peerj.10098

Justine, J.-L., Gey, D., Vasseur, J., Thévenot, J., Coulis, M. & Winsor, L. (2021) Presence of the invasive land flatworm Platydemus manokwari (Platyhelminthes, Geoplanidae) in Guadeloupe, Martinique and Saint Martin (French West Indies). Zootaxa, 4951, 381–390. http://doi.org/10.11646/zootaxa.4951.2.11

Justine, J.-L., Winsor, L., Barrière, P., Fanai, C., Gey, D., Han, A. W. K., et al. (2015) The invasive land planarian Platydemus manokwari (Platyhelminthes, Geoplanidae): records from six new localities, including the first in the USA. PeerJ, 3, e1037. http://doi.org/10.7717/peerj.1037

Justine, J.-L., Winsor, L., Gey, D., Gros, P. & Thévenot, J. (2014) The invasive New Guinea flatworm Platydemus manokwari in France, the first record for Europe: time for action is now. PeerJ, 2, e297. http://doi.org/10.7717/peerj.297

Justine, J.-L., Winsor, L., Gey, D., Gros, P. & Thévenot, J. (2018) Giant worms chez moi! Hammerhead flatworms (Platyhelminthes, Geoplanidae, Bipalium spp., Diversibipalium spp.) in metropolitan France and overseas French territories. PeerJ, 6, e4672. http://doi.org/10.7717/peerj.4672

Justine, J.-L., Winsor, L., Gey, D., Gros, P. & Thévenot, J. (2020b) Obama chez moi! The invasion of metropolitan France by the land planarian Obama nungara (Platyhelminthes, Geoplanidae). PeerJ, 8, e8385. http://doi.org/10.7717/peerj.8385

Justine, J. L. & Winsor, L. (2020) First record of presence of the invasive land flatworm Platydemus manokwari (Platyhelminthes, Geoplanidae) in Guadeloupe. Preprints, 2020, 2020050023. http://doi.org/10.20944/preprints202005.0023.v1

Kass, J. M., Muscarella, R., Galante, P. J., Bohl, C. L., Pinilla Buitrago, G. E., Boria, R. A., Soley Guardia, M. & Anderson, R. P. (2021) ENMeval 2.0: redesigned for customizable and reproducible modeling of species’ niches and distributions. Methods in Ecology and Evolution. http://doi.org/10.1111/2041-210X.13628

Kumar, S., Stecher, G., Li, M., Knyaz, C. & Tamura, K. (2018) MEGA X: molecular evolutionary genetics analysis across computing platforms. Molecular Biology and Evolution, 35, 1547. http://doi.org/10.1093/molbev/msy096

Lago-Barcia, D., González-López, J. R. & Fernández-Álvarez, F. Á. (2020) The invasive land flatworm Obama nungara (Platyhelminthes: Geoplanidae) reaches a natural environment in the oceanic island of São Miguel (Açores). Zootaxa, 4830, 197–200.

Leigh, J. W. & Bryant, D. (2015) POPART: full-feature software for haplotype network construction. Methods in Ecology and Evolution, 6, 1110–1116. http://doi.org/10.1111/2041-210x.12410

Lowe, S., Browne, M., Boudjelas, S. & De Poorter, M. (2000) 100 of the World’s Worst Invasive Alien Species. A selection from the Global Invasive Species Database. Published by The Invasive Species Specialist Group (ISSG) a specialist group of the Species Survival Commission (SSC) of the World Conservation Union (IUCN), 12pp. First published as special lift-out in Aliens 12, December 2000. Updated and reprinted version: November 2004.

Mori, E., Giulia, M., Panella, M., Montagna, M., Winsor, L., Justine, J.-L., Menchetti, M., Schifani, E., Melone, B. & Mazza, G. (2022) Opening Pandora’s box: the invasion of alien flatworms in Italy. Biological Invasions, 24, 205–216. http://doi.org/10.1007/s10530-021-02638-w

Murchie, A. K. & Justine, J.-L. (2021) The threat posed by invasive alien flatworms to EU agriculture and the potential for phytosanitary measures to prevent importation. Technical note prepared by IUCN for the European Commission.

Negrete, L., Lenguas Francavilla, M., Damborenea, C. & Brusa, F. (2020) Trying to take over the world: Potential distribution of Obama nungara (Platyhelminthes: Geoplanidae), the Neotropical land planarian that has reached Europe. Global Change Biology, 26, 4907–4918. http://doi.org/10.1111/gcb.15208

Phillips, S. J., Anderson, R. P., Dudík, M., Schapire, R. E. & Blair, M. E. (2017) Opening the black box: An open source release of Maxent. Ecography, 40, 887–893. http://doi.org/10.1111/ecog.03049

Quinteiro, J., Rodríguez Castro, J., Castillejo, J., Iglesias Piñeiro, J. & Rey Méndez, M. (2005) Phylogeny of slug species of the genus Arion: evidence of monophyly of Iberian endemics and of the existence of relict species in Pyrenean refuges. Journal of Zoological Systematics and Evolutionary Research, 43, 139–148. http://doi.org/10.1111/j.1439-0469.2005.00307.x

Rowson, B., Anderson, R., Turner, J. A. & Symondson, W. O. (2014) The slugs of Britain and Ireland: undetected and undescribed species increase a well-studied, economically important fauna by more than 20%. PLoS ONE, 9, e91907. http://doi.org/10.1371/journal.pone.0091907

Sluys, R. (2016) Invasion of the Flatworms. American Scientist, 104, 288–295.

Solà, E., Álvarez-Presas, M., Frías-López, C., Littlewood, D. T. J., Rozas, J. & Riutort, M. (2015) Evolutionary analysis of mitogenomes from parasitic and free-living flatworms. PLoS ONE, 10, e0120081. http://doi.org/10.1371/journal.pone.0120081

Winsor, L. (1983) A revision of the Cosmopolitan land planarian Bipalium kewense Moseley, 1878 (Turbellaria: Tricladida: Terricola). Zoological Journal of the Linnean Society, 79, 61–100. http://doi.org/10.1111/j.1096-3642.1983.tb01161.x

